# Historical genomes reveal scale-dependent predictability of climate adaptation

**DOI:** 10.64898/2026.08.18.744306

**Authors:** Paul Battlay, Saila Kabir, Vanessa C. Bieker, Sarah L. F. Martin, Vid Terzer, Loren H Rieseberg, Keyne Monro, Alexandre Fournier-Level, John R. Stinchcombe, Michael D. Martin, Kathryn A. Hodgins

**Author notes:** Authors contributed equally (Paul Battlay, Saila Kabir).

## Abstract

Predicting evolution remains a central challenge in biology. Contemporary spatial patterns are increasingly used as space-for-time proxies to forecast evolutionary responses to environmental change, yet the reliability of such predictions—and whether it varies among biological scales—remains unclear. Using historical and contemporary genomes of the invasive weed *Ambrosia artemisiifolia*—spanning the species’ native range, invasions on two continents and nearly two centuries—we show that genomically predicted flowering-time clines remained stable in the native range while introduced populations re-evolved them. Haploblocks—likely structural variants—were likewise temporally stable in the native range, and two showed striking parallel evolution across all three ranges. Climate-associated SNPs with the strongest contemporary clines showed the greatest temporal change, with limited and variable parallelism among introduced ranges. Together, these results suggest adaptive evolution is partly predictable even when individual genomic trajectories remain flexible and contingent, with predictability emerging most clearly at the level of polygenic traits and large structural variants.

## Introduction

Predicting evolutionary responses to human-driven environmental change is one of the most pressing challenges in evolutionary biology. Parallel evolution across replicated environmental gradients suggests that adaptation can be repeatable, raising the possibility that contemporary spatial patterns may help forecast future adaptation^1^. However, historical contingency, demographic history, and genomic architecture can all limit evolutionary repeatability^2–4^, leaving unresolved the question of whether adaptive responses are sufficiently repeatable to be predictable.

Much of our understanding of climate adaptation comes from spatial patterns across environmental gradients, including trait variation along clines and genotype–environment associations^5,6^. Genomic patterns are increasingly used to infer vulnerability to future climate change and the potential for evolutionary rescue^7,8^. These space-for-time approaches assume that contemporary spatial relationships between climate and genomic variation are informative about evolutionary change through time^9^. This assumption may fail when recent environmental change or colonisation leaves populations out of equilibrium, with demographic history and genomic architecture further shaping the variation available to selection and its subsequent trajectory. Recent work using preserved samples has begun to validate climate-associated genomic change directly (e.g., ^10^) but the extent to which spatial clines reliably predict temporal evolutionary trajectories remains largely untested.

Sequencing archived historical specimens now allows direct observation of adaptive genomic change^11–13^. Biological invasions provide particularly powerful systems in this regard, generating replicated natural experiments in which populations with shared ancestry adapt independently to similar climatic gradients across multiple introduced ranges^14,15^. If contemporary spatial clines reflect stable adaptive relationships, existing native-range clines may remain relatively consistent through time, while clines in introduced populations should progressively evolve following introduction. Alternatively, demographic history, genomic architecture or shifting selective environments may cause temporal instability in the native range or divergence in adaptive patterns across ranges. Evolutionary repeatability, and hence predictability, may also differ among levels of genetic organisation. Similar polygenic responses can arise through different contributing variants^3^, whereas haploblocks—large linked genomic regions—may be repeatedly recruited if they preserve combinations of adaptive alleles or exert larger effects than individual SNPs^16,17^.

Common ragweed (*Ambrosia artemisiifolia*) is a globally invasive annual plant native to North America that was introduced into Europe in the mid-1800s and into Australia in the early 1900s^18^. European populations experienced multiple introductions and extensive admixture with little evidence of a bottleneck, whereas the Australian invasion underwent a substantial population bottleneck and was founded from distinct native-range sources^18^. Despite these contrasts, both invasions rapidly expanded across climatic gradients and repeatedly established latitudinal clines in flowering time, paralleled by genomic signatures of climate adaptation ^18–22^. Extensive herbarium collections spanning both native and introduced ranges provide a unique opportunity to test whether contemporary spatial patterns of adaptation predict temporal evolutionary change^13,23,24^.

Using historical and modern genomic data spanning nearly two centuries, we tested whether contemporary spatial patterns of adaptation predict temporal evolutionary change. We predicted that climate-associated loci—and particularly those forming the strongest contemporary clines— would show greater temporal change than matched genomic controls, with the greatest responses occurring where introduced populations experienced the strongest climatic mismatch. We further tested whether adaptive clines remained stable in the native range and were repeatedly re-established across independent invasions, and whether this repeatability differed among individual SNPs, flowering-time polygenic scores and haploblocks.

## Results

### Climate-associated loci show greater temporal change

The core assumption of space-for-time approaches is that contemporary climate associations in space predict evolutionary change through time. To test this, we combined 90 newly sequenced herbarium specimens, additional coverage sequencing of existing herbarium samples, and previously published modern and historical genomes^11,18^(Table S1; Table S2), and called SNPs across 769 samples of common ragweed spanning its native North American range and invasions in Europe and Australia, collected over ∼180 years. We then identified candidate SNPs associated with four minimally correlated bioclimatic variables (BIO1, mean annual temperature; BIO2, mean diurnal temperature range; BIO12, annual precipitation; and BIO15, precipitation seasonality), yielding between 939 and 3,206 candidate loci per climate variable (Table S3).

Candidate loci associated with contemporary adaptation exhibited larger temporal shifts in allele–environment clines than allele frequency–matched null SNPs (all Wilcoxon test *p*-values <= 1.37×10^-12^; Fig. 1A). This pattern was consistent across all three ranges and all four climate variables, indicating that temporal change occurs specifically at loci associated with climate, rather than reflecting genome-wide shifts. Across all climate variables, candidate loci in both European and Australian ranges exhibited significantly larger temporal cline shifts than candidates from the native North American range (Wilcoxon tests on null-relative *Z*-scores; Fig. 1B), except for Australian BIO15 candidates, which showed significantly smaller shifts.

**Figure 1.**
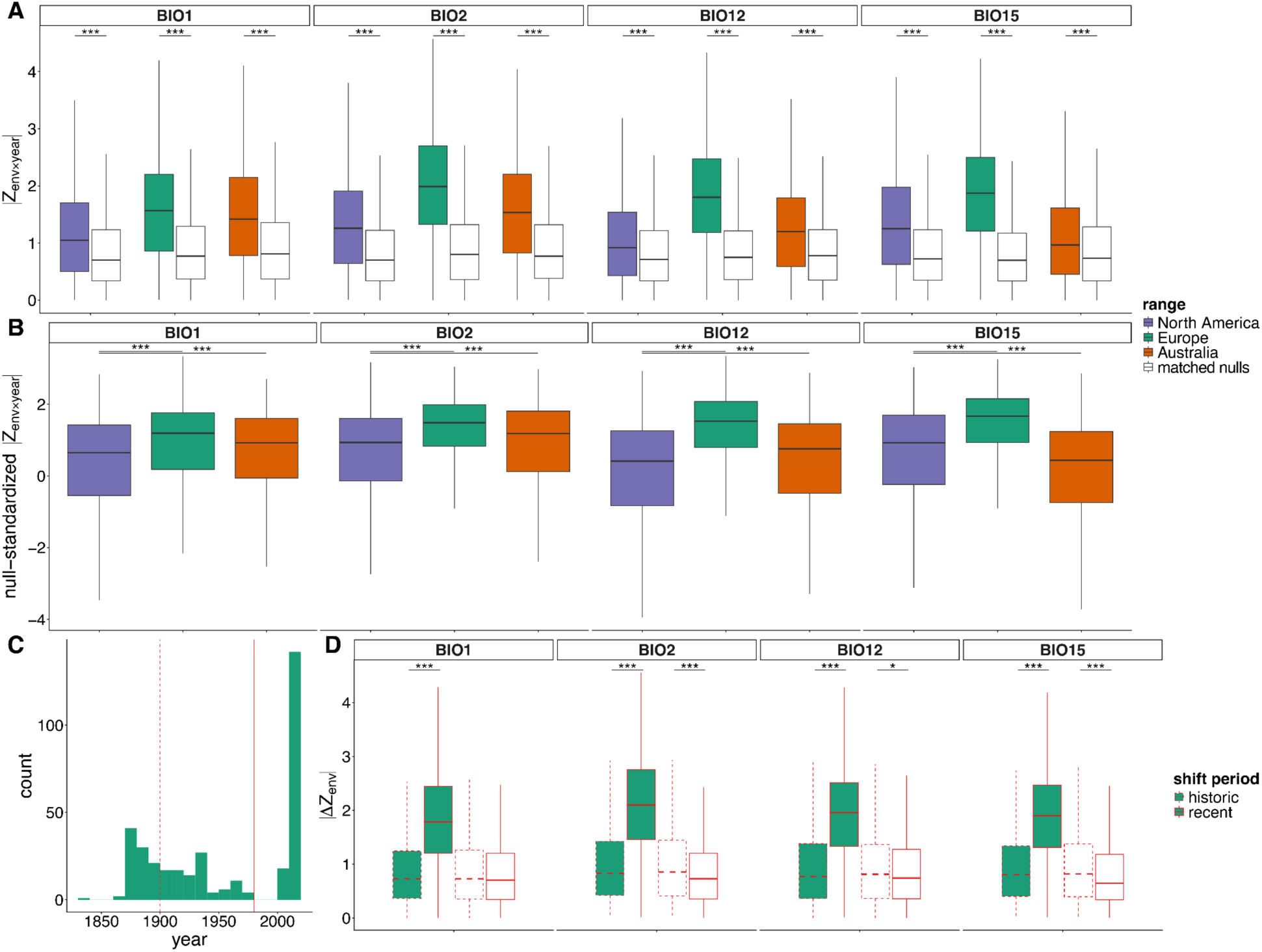
Temporal cline shifts in range-specific candidate loci. Boxes show medians and interquartile ranges; whiskers indicate 1.5×IQR. Asterisks indicate significance of differences between groups in Wilcoxon tests (\**p* < 0.05; \*\**p* < 0.01; \*\*\**p* < 0.001). **A.** Distribution of temporal allele-frequency shifts at candidate loci (coloured boxes) compared with null loci (white boxes), shown for each environmental variable (BIO1, BIO2, BIO12, BIO15) and geographic range (North America, Europe, Australia). Shifts are quantified as the magnitude of the environment × year interaction term divided by its standard error from the genotype–environment–time model (*|Z_env×year_|*). **B.** Candidate SNP shifts standardized relative to the range-specific distribution of null SNPs. For each candidate, the *|Z_env×year_|* statistic was centred on the median of allele-frequency matched null SNPs and scaled by their median absolute deviation (MAD), yielding a null-relative standardized statistic. Positive values indicate candidate loci exhibiting larger temporal cline shifts than expected under the genome-wide null distribution. **C.** Distribution of European sampling dates used in the three-category time model (pre-1900, *n* = 97; 1900–1980, *n* = 109; post-1980, *n* = 160). Vertical dashed lines indicate the boundaries between sampling periods. **D.** Magnitude of temporal change in allele–environment clines between sampling periods (historic: pre-1900 vs. 1900–1980; recent: 1900–1980 vs. post-1980) in European candidates (green boxes) compared to allele frequency-matched null SNPs (white boxes), measured as the absolute difference in the standardized environmental effect size between time intervals (*|ΔZ_env_|*).

Modelling three temporal bins in Europe (where herbarium sampling provided sufficient temporal resolution; Fig. 1C), we asked whether cline evolution intensified in particular periods of the invasion. Candidate SNPs exhibited significantly larger cline shifts in the more recent interval (1900–1980 vs. modern) than in the earlier historic interval (<1900 vs. 1900–1980) for all environmental variables (Fig. 1D), indicating that much of the observed clinal evolution occurred during the later stages of invasion. Matched null SNPs, in contrast, showed no comparable acceleration (Fig. 1D).

### Climatic mismatch and temporal allele-frequency change are asymmetrically distributed across environmental gradients

Invasive species are often introduced to climates that are different from their native source. If adaptive change acts to reduce this ‘climatic mismatch’, the frequency of climate-associated variants should show a temporal shift in introduced populations as they adapt, with larger shifts expected in regions where climatic mismatch is greater. To test this, we quantified climatic mismatch between introduced populations and their inferred source, and asked whether temporal allele-frequency shifts of climate candidates were correspondingly asymmetric across environmental gradients. Locator^25^ predicted geographically distinct source regions for the two invaded ranges (Fig. 2A). European samples were most frequently assigned to northern and eastern portions of the native North American range, whereas Australian samples were assigned to more southern regions.

**Figure 2.**
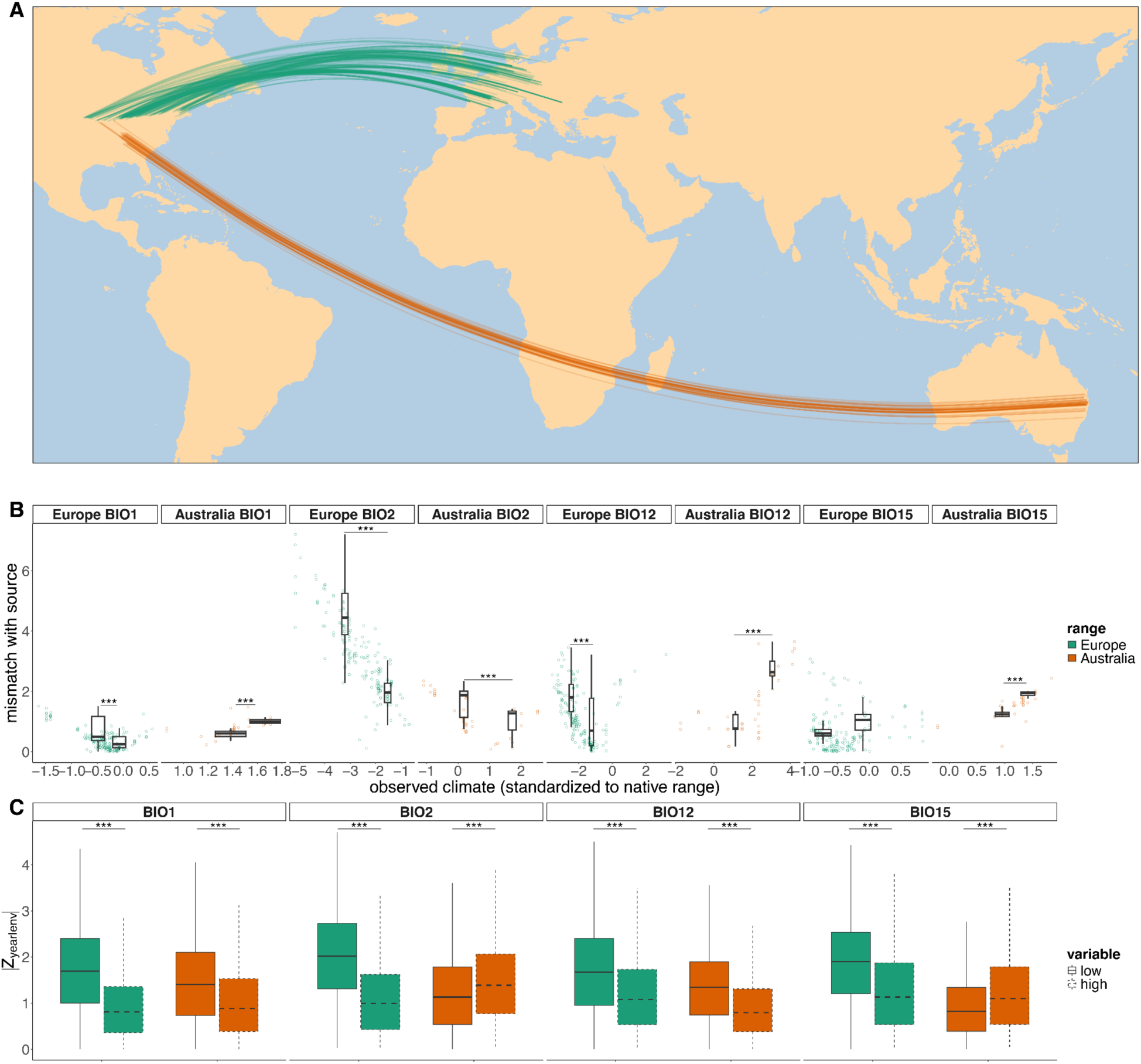
Environmental mismatch and asymmetric temporal allele-frequency change across invaded ranges. **A.** Locator predictions for historic samples from Europe and Australia based on models trained on historic North American samples. Lines connect sampling locations to their inferred source locations in the native range. **B.** Relationship between standardized climate at the sample location (*x*-axis; scaled relative to the climate distribution of North American training samples) and the mismatch between the sample location and inferred source location (*y*- axis). Boxes compare mismatch at the 25th and 75th percentile. **C.** Distribution of temporal allele-frequency change at candidate loci, quantified as the slope of allele frequency with respect to year, evaluated at the low (10th percentile) and high (90th percentile) values of each environmental gradient. Values are shown as standardized effect sizes (slope/SE; *|Z_year|env_|*). Boxes show medians and interquartile ranges; whiskers indicate 1.5×IQR. Asterisks indicate significance of differences between groups in Wilcoxon tests (\**p* < 0.05; \*\**p* < 0.01; \*\*\**p* < 0.001).

Introduced samples frequently occupied climates that differed from those of their inferred native source, and the magnitude of this climatic mismatch varied along environmental gradients (Fig. 2B). Mismatch differed significantly between upper and lower quartiles for BIO1, BIO2, and BIO12 in both Europe and Australia (all *p* < 0.001), and for BIO15 in Australia but not Europe. These relationships indicate that populations at one cline end experience substantially greater climatic mismatch relative to their inferred source. We next asked whether temporal change was similarly concentrated toward particular portions of environmental gradients. Across all variables in invaded ranges, temporal allele-frequency change at candidate loci differed significantly between the two extremes of environmental gradients (Fig. 2C; *p* < 0.001). These asymmetries in temporal change were also significantly greater than those observed in a matched set of nulls (Fig. S1). This result was robust to uncertainty in Locator source assignment, with all range-by-variable estimates retaining the same direction in at least 98.8% of 1,000 genomic-window bootstrap replicates and all 95% bootstrap intervals excluding zero.

Despite widespread climatic mismatch relative to inferred source regions, and strong asymmetry in temporal allele-frequency change, the correspondence between mismatch and cline-end shifts differed between invaded ranges. In Europe, for variables showing strong mismatch (BIO1, BIO2, BIO12), the cline end with the greatest climatic mismatch also showed the largest temporal allele-frequency shifts. Contrastingly, in Australia, climate mismatch and cline-end allele-frequency shifts only corresponded for BIO15. In the native North American range, cline- end asymmetry was much weaker (Fig. S1), with significant differences only for BIO12.

### Strong contemporary climate clines exhibit elevated historical temporal change

Theory predicts that stronger environmental selection will generate steeper clines for a given rate of dispersal^26^. Accordingly, if the steepness of clines reflects the strength of selection, loci with the strongest climate associations should exhibit the greatest evolutionary responses to environmental change over time. We therefore tested whether cline shifts over time were correlated with the strength of modern allele frequency clines. Across all ranges and climate variables, candidate SNPs showed a positive relationship between contemporary cline steepness and historical temporal change (Fig. 3). SNPs exhibiting the strongest contemporary clines tended to show the greatest temporal shifts across all four climatic variables. Matched null SNPs generally exhibited weaker relationships, suggesting that the association was enriched among climate-associated loci rather than reflecting genome-wide patterns of spatial and temporal allele-frequency change. These results suggest that loci contributing most strongly to contemporary climate adaptation across space tend to experience elevated evolutionary change over time.

**Figure 3.**
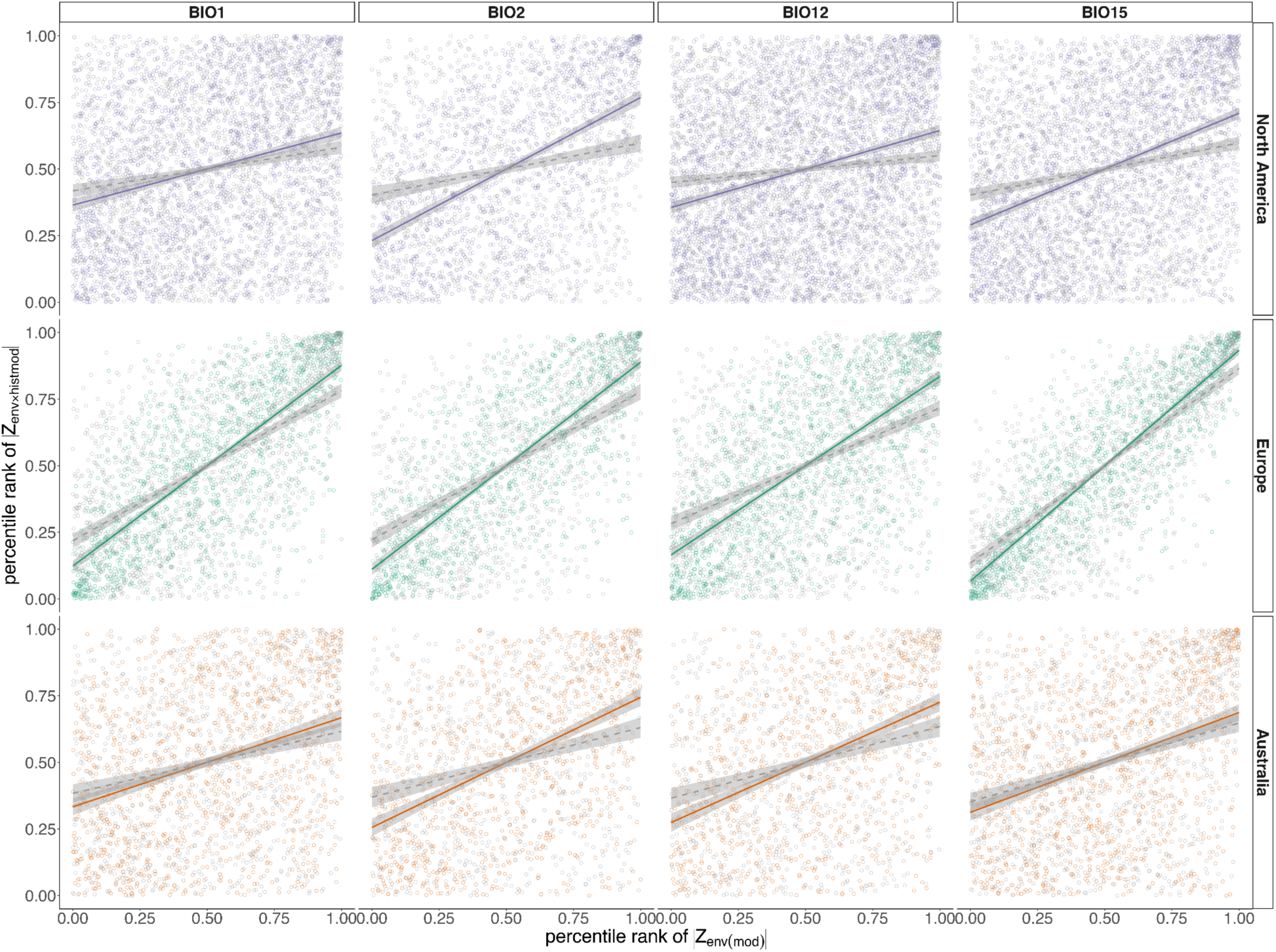
Relationship between environmental effect size and temporal cline shift. Percentile rank of temporal cline shift (|*Z_env×histmod_*|), estimated from the contrast between modern and historic environmental clines, plotted against percentile rank of environmental effect size (|*Z_env(mod)_*|), estimated from the modern environmental cline. Results are shown for each geographic range and climate variable. Candidate SNPs are shown in colour and matched null SNPs in grey. Lines show linear model fits where significant (p < 0.05), with solid lines for candidate SNPs and dotted lines for null SNPs.

### Signatures of parallel cline evolution across ranges

If adaptive clines are repeatedly reestablished during invasion, loci associated with climate adaptation in North America should show parallel clines in modern invaded ranges. To test this, we asked whether loci associated with climate in North America also showed stronger clines than matched null SNPs in the invaded ranges (Fig. 4A). In Europe, North American candidate loci showed slightly stronger clines than nulls for BIO1 and BIO15, but not for BIO2 or BIO12. In Australia, candidate loci showed stronger modern clines for BIO15, weaker clines for BIO1, and clines no different from nulls for BIO2 or BIO12. We further asked whether invaded-range clines matched the direction of the modern North American cline, and how this changed over the course of invasions (Fig. 4B). For some range–environment combinations, North American candidate loci showed increasing directional concordance through time, whereas others showed divergence from native-range cline direction. In Europe, concordance increased over time for BIO1 and BIO12, whereas BIO15 diverged from the native cline direction. In Australia, concordance increased for BIO15 but declined for BIO12. Significant patterns were absent in null SNPs (Fig. S2). Together, these results suggest that loci associated with native-range climate adaptation remain evolutionarily dynamic during invasion, but parallel climate adaptation involves only partial and variable reuse of native-range clinal patterns. This variable pattern of parallelism is consistent with previous findings in common ragweed showing only partial overlap (∼14–24%) in climate associations between native and invaded ranges^18,22^.

**Figure 4.**
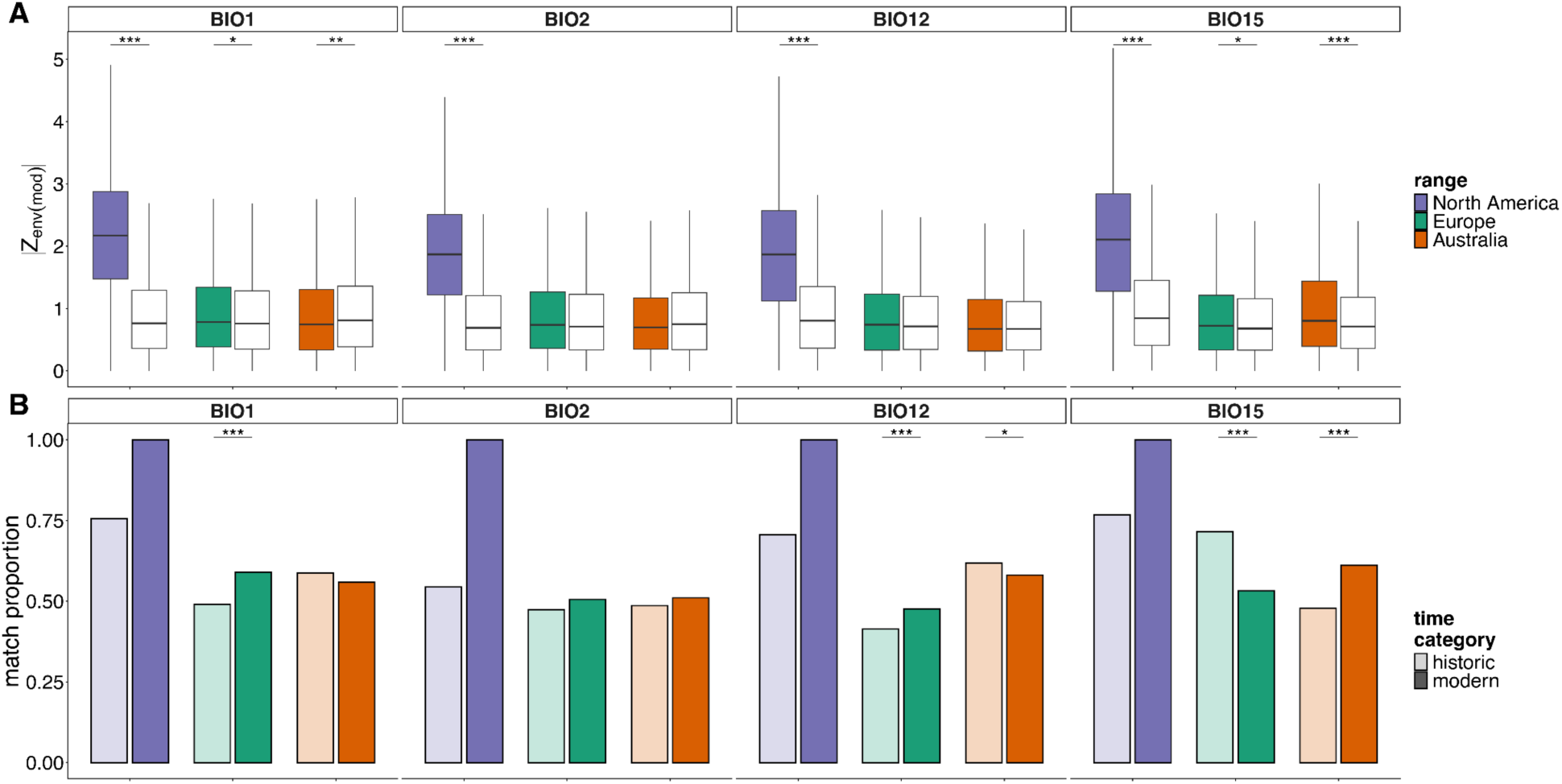
Native-invaded parallelism in climate adaptation. **A.** Distribution of modern cline slope magnitudes for North American candidate loci (coloured boxes) in each range compared with matched null loci (white boxes). Shifts are measured as the magnitude of the modern estimate of the environmental effect size from the genotype–environment–time model. Boxes show medians and interquartile ranges; whiskers indicate 1.5×IQR and asterisks indicate significance of differences between groups in Wilcoxon tests (\**p* < 0.05; \*\**p* < 0.01; \*\*\**p* < 0.001). **B.** Directional alignment with native-range clines. Bars indicate the proportion of North American candidate loci whose clines match the direction of the modern North American cline, shown separately for historic (pre-1980) and modern samples. Asterisks denote significant changes in matching frequencies between time categories based on McNemar’s tests (\**p* < 0.05; \*\**p* < 0.01; \*\*\**p* < 0.001). North American data are included for comparison, but McNemar’s tests were not applicable because all loci matched the modern native cline by definition, resulting in degenerate contingency tables with no discordant pairs.

### Flowering-time clines are stable in the native range and re-evolve during invasion

Having characterised temporal change at individual climate-associated SNPs, we next asked whether predictability differs for a polygenic trait. We expected that adaptive latitudinal clines in flowering time would remain stable through time in the native range, and evolve in parallel following introduction. To test this, we derived polygenic scores (PGS) from genome-wide associations (93 LD-clumped SNPs; GWAS *p* < 1×10^-5^) for flowering onset from a subset of modern samples (*n* = 212). We then predicted flowering onset across all modern and historic samples and tested for latitudinal clines across time and space. In the native North American range, flowering onset PGS exhibited a strong and consistent latitudinal cline, with higher-latitude populations predicted to flower earlier, and no evidence for temporal change in clines between historic and modern samples (Fig. 5). In contrast, European samples showed no historic cline, but a strong negative relationship in the modern time period, indicating the rapid evolution of a latitudinal flowering-time gradient following introduction to Europe. Australian populations showed weaker patterns overall, but effect sizes were consistent with an emerging cline: historic samples showed little to no latitudinal structure, whereas modern samples exhibited a marginally negative cline with a significant interaction between latitude and time. These patterns were qualitatively robust both to changes in GWAS significance thresholds (Fig. S3) and excluding predictions for individuals with phenotypes (those used to train the PGS; Fig. S4). These patterns suggest that genetic clines in flowering-time can evolve in parallel across invaded ranges, recapitulating native-range patterns through time.

**Figure 5.**
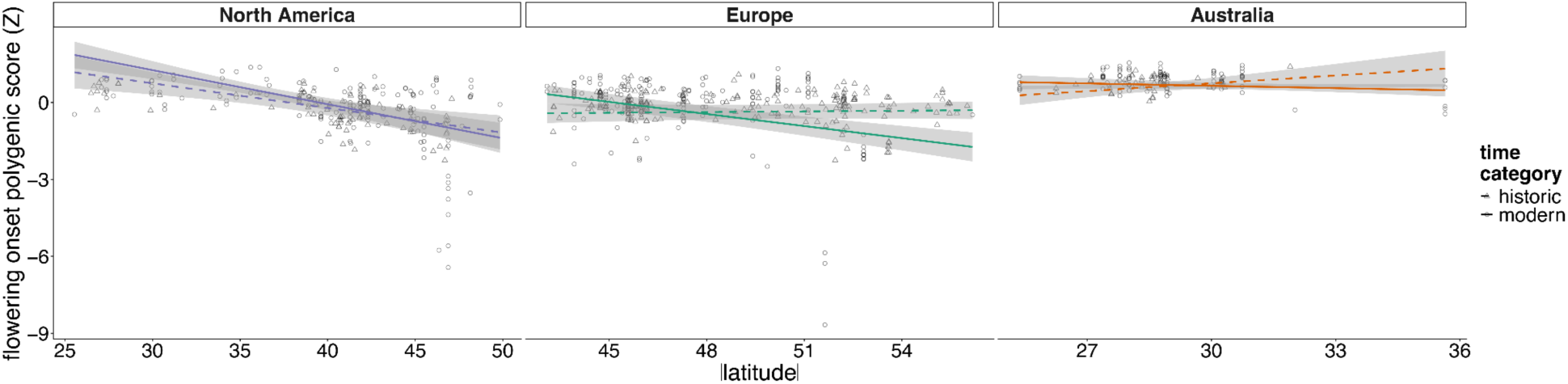
Latitudinal clines in flowering onset polygenic scores across native and invaded ranges. Polygenic scores (PGS; standardized) for flowering onset plotted against absolute latitude for each range. Points represent individual samples (triangle = historic; circle = modern). Lines (dashed = historic; solid = modern) show fitted relationships from linear models including latitude, time (historic vs modern), and their interaction (with the first two principal components of genome-wide SNP variation as covariates); shaded regions indicate 95% confidence intervals.

### Climate-associated haploblocks are stable in the native range and show parallel cline evolution across invasions

Having found that a polygenic trait re-evolved clinal structure following invasion, we next asked whether large haploblocks—population-genomic signatures of inversions or other sources of recombination suppression—show the same temporal signature. Previous work in ragweed has shown that large haploblocks contribute disproportionately to climate associations and adaptive traits^18^. To recover these regions from our temporally resolved SNP data, we scanned the genome for distortions in local population structure^27^, identifying 34 haploblocks with a global minor allele frequency > 0.05 (Table S4). Of these, 29 corresponded to haploblocks identified previously^18^, and 14 overlapped known inversion polymorphisms in the phased diploid reference. We fitted environmental association models to each haploblock’s genotypes using genotype–environment–time models. More than half of the haploblocks (20/34) showed evidence of climate association in North America, with effects exceeding the 95th percentile of a null SNP distribution for at least one climate variable (Fig. 6A). These candidate haploblocks were stable in North America, with no changes in cline direction between historic and modern samples (Fig. 6B). In contrast, haploblocks in invaded ranges often showed increasing directional concordance with native-range clines through time. The proportion of haploblocks whose clines matched the direction of the native range increased over time for annual mean temperature (BIO1) and annual precipitation (BIO12) in both Europe and Australia. For precipitation seasonality (BIO15), concordance increased in Europe but declined markedly in Australia. Two haploblocks showed particularly striking evidence of parallel evolution: in both invasions their clines shifted through time toward the native-range direction (Fig. 6C). These genomic region-level patterns, like flowering-time polygenic scores, were more consistently parallel across invasions than the underlying individual SNPs—indicating that the predictability of climate adaptation depends on the genomic scale at which it is measured.

**Figure 6.**
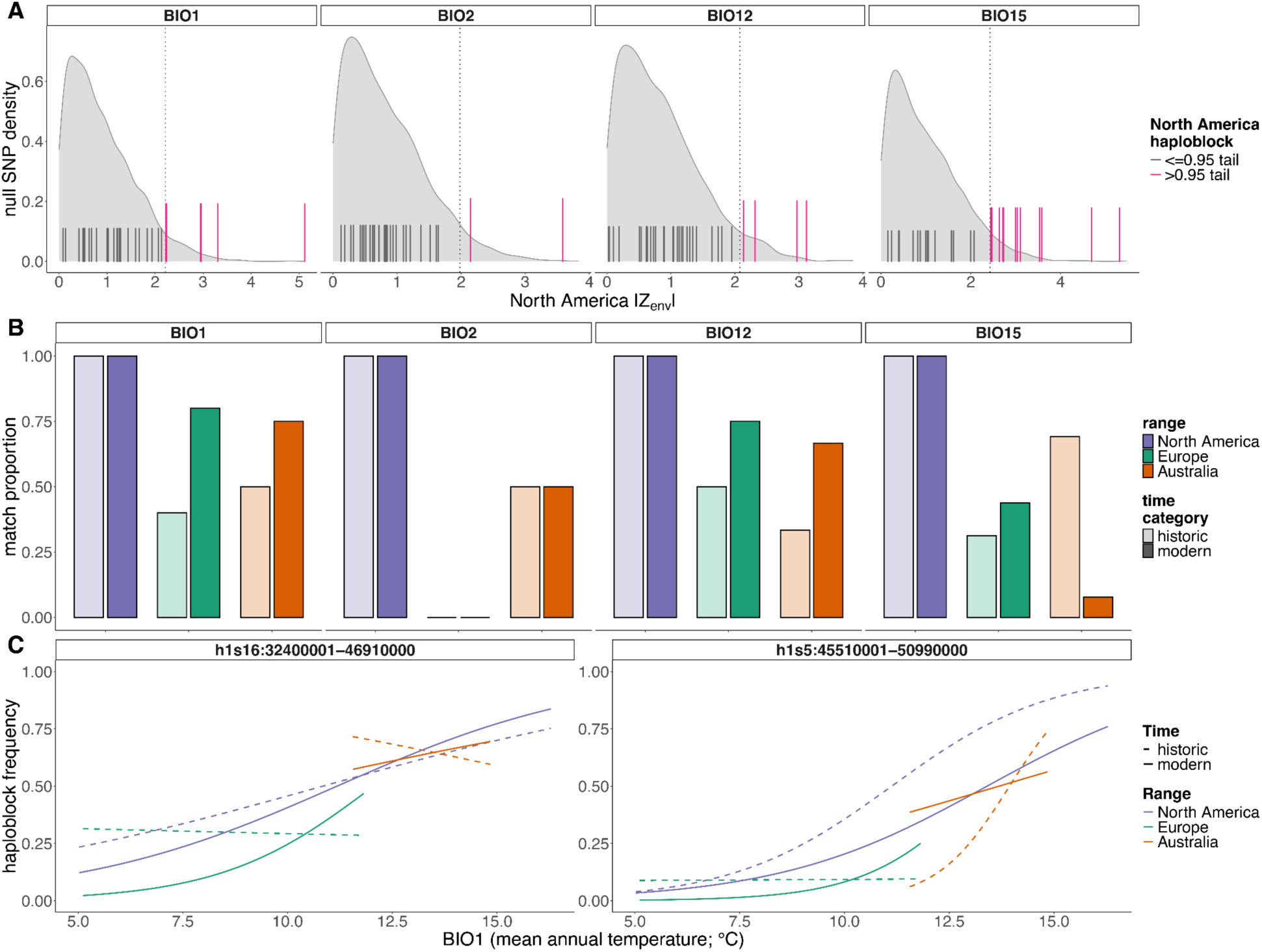
Climate-associated haploblocks show enrichment, temporal stability, and parallel cline evolution. **A.** Distribution of environmental association statistics (|Z_env|) for haploblock allele frequency–matched null SNPs in North America for each climate variable (BIO1, BIO2, BIO12, BIO15). Solid vertical lines indicate haploblock values, with pink lines denoting haploblocks exceeding the 95th percentile of the null distribution (represented by the dotted line) and grey lines indicating those below this threshold. **B.** Proportion of haploblocks whose clines match the direction of the North American cline in each range, shown separately for historic (light shading) and modern (dark shading) samples. North American haploblocks are uniformly stable over time, whereas concordance in invaded ranges varies among climate variables and increases over time for BIO1 and BIO12. **C.** Two haploblocks showing parallel clines across all three ranges for BIO1. Lines show fitted clines from betabinomial models, with historic (dashed) and modern (solid) timepoints.

## Discussion

Genomic offset and related space-for-time frameworks increasingly use contemporary patterns of adaptation to infer vulnerability to environmental change^7,9^. In principle, patterns of adaptation across space should predict evolutionary response over time, yet this central assumption has rarely been tested directly (but see ^28^). By comparing historical and contemporary genomes from the native range and two replicated invasions, we show that contemporary climate-associated clines most consistently predicted temporal evolutionary change within ranges, whereas parallelism across invasions was more variable and emerged more clearly for flowering-time polygenic scores and large haploblocks than at individual SNPs.

At the SNP level, these results provide direct empirical support for a central assumption of space-for-time inference: contemporary climate associations contain information about evolutionary change through time. Contemporary climate adaptation candidate loci consistently exhibited greater temporal change than nulls. Moreover, loci with the strongest contemporary clines generally exhibited the greatest temporal shifts through time, and these relationships were consistently stronger than those observed for nulls. This suggests that loci with the largest effects on contemporary climate-associated differentiation tend to respond more strongly over short evolutionary timescales.

Although both invaded ranges evolved climate-associated clines through time, the degree of evolutionary predictability differed substantially between Europe and Australia. In Europe, the cline ends experiencing the greatest climatic mismatch also underwent the largest temporal allele-frequency shifts, and directional concordance with native-range clines increased through time for most variables, consistent with relatively predictable adaptation following invasion. In contrast, temporal cline shifts in Australian populations were less predictable, showing weaker association with climatic mismatch and weaker convergence toward native-range clines. These differences are consistent with the variation in genetic architecture between ranges^4^, and may also reflect distinct demographic histories (including a more recent introduction, stronger bottlenecks and more restricted source ancestry in Australia^18^) or divergent selective regimes between continents. These results indicate that climatic mismatch alone may not consistently predict adaptive change, as its effects may depend on demographic history, genomic architecture and broader selective context.

Flowering-time polygenic score clines provided one of the clearest examples of parallel adaptive evolution through time: latitudinal clines remained stable across nearly two centuries in native North American populations, but were initially weak or absent in introduced populations, and re-evolved over the course of invasions. This extends prior evidence of parallel flowering-time clines in ragweed^19,21^ and other invasive plants^29^. Similar genomic scores have captured century-long trait change in *Arabidopsis thaliana*^30^, here applied across replicated invasions. This repeated emergence, despite differing demographic histories and partial reuse of climate-associated loci, illustrates how concordance at individual variants can underestimate repeatability at the phenotypic level: populations can converge on similar trait optima through different combinations of allele-frequency shifts because genetic redundancy and population-specific constraints and biases provide alternative routes to adaptation^3^.

We also observed repeatability at the level of large haploblocks. Climate-associated haploblocks maintained consistent directions of environmental association across nearly two centuries in North America, and many showed increasing concordance with native-range clines across invaded ranges through time—consistent with large structural variants contributing disproportionately to local climate adaptation by holding favourable combinations of alleles together against recombination^16,31^. Overall, our results suggest that evolutionary predictability was generally greater at the level of traits and major haploblocks, and least clearly at individual variants—the scale on which space-for-time forecasts most often rely.

Beyond variation across ranges and genomic scales, the tempo of adaptation was itself uneven through time. In Europe, where herbarium sampling resolved multiple historical intervals, climate-associated loci showed substantially stronger cline shifts in the recent interval (1900–1980 vs. modern) than earlier in the invasion. This acceleration may be demographic: European ragweed remained localized until major range expansion in the mid-twentieth century^32,33^, so early populations were likely too small and fragmented for efficient selection, with coordinated clinal evolution emerging only as populations expanded. These recent shifts could partly reflect ascertainment bias, since candidates were identified in the modern period. However, the greater temporal stability of candidates in the native range—where the same identification framework was applied—and the absence of comparable shifts in matched null SNPs argue against a purely methodological explanation. Such rapid recent evolution suggests that invasive populations can respond quickly to changing climates once they become widespread, potentially facilitating further expansion and complicating management.

Together, our results suggest that improving space-for-time forecasts requires identifying the levels of biological organisation at which adaptive responses remain repeatable through time and across geographic ranges—a challenge also highlighted by regional variation in adaptive clines and their genomic architectures in other plant species^34,35^. In common ragweed, repeatability was most evident for flowering-time polygenic scores and large haploblocks, and weakest for individual variants. Because space-for-time forecasts typically rest on SNP-level structure, they may rely on precisely the least repeatable level of adaptive change. Resolving these dynamics directly from historical genomes reveals both the promise and the limits of using present-day spatial variation to anticipate evolution under rapid climate change.

## Acknowledgements

We thank the curators of the following herbaria for allowing us to destructively sample their collections: AD, AK, BRI, CANB, CHR, HO, MEL, NSW, and PERTH. We thank Juan Francisco Esteves Ramírez and Paula Kosel for assistance with the laboratory work. Computational analyses were performed on the Monash MASSIVE M3 and Digital Research Alliance of Canada high-performance computing platforms. This work was supported by an Australian Research Council Discovery Project grant DP220102362 awarded to KAH, MDM, JRS and AFL.

## Author contributions

**Paul Battlay\*** Methodology, Software, Formal analysis, Investigation, Data Curation, Writing - Original Draft, Writing - Review & Editing, Visualization, Supervision

**Saila Kabir\*** Methodology, Software, Formal analysis, Investigation, Writing - Original Draft, Writing - Review & Editing, Visualization

**Vanessa C. Bieker** Investigation, Writing - Review & Editing

**Sarah L. F. Martin** Investigation

**Vid Terzer** Investigation

**Loren H Rieseberg** Resources, Writing - Review & Editing

**Keyne Monro** Conceptualization, Methodology, Software, Formal analysis, Writing - Review & Editing, Supervision

**Alexandre Fournier-Level** Writing - Review & Editing, Funding acquisition

**John R. Stinchcombe** Writing - Review & Editing, Funding acquisition

**Michael D. Martin** Investigation, Resources, Writing - Review & Editing, Funding acquisition

**Kathryn A. Hodgins** Conceptualization, Methodology, Software, Writing - Original Draft, Writing

- Review & Editing, Supervision, Project administration, Funding acquisition

*Authors contributed equally

## Data availability

The phased diploid reference genome assembly used in this study is available from NCBI under BioProject IDs PRJNA929657 and PRJNA929658, organelle assemblies are available from FigShare (doi.org/10.26180/33201693). Individual sample resequencing data are available from ENA under BioProject IDs PRJEB48563, PRJNA339123, and PRJEB34825, and from SRA under BioProject IDs PRJNA1139307 and PRJNA1497700. Phenotypic data are available from github.com/lotteanna/traitclines. RepAdapt pipeline code used to produce alignments is available from github.com/JimWhiting91/RepAdapt/tree/main/snp_calling_pipeline/mpileup_pipeline. Code for analyses is available from github.com/pbattlay/ragweed-space-and-time.

## Online methods

### Reference genome

This study used the primary haplotype of the diploid *Ambrosia artemisiifolia* reference genome described in^22^. To exclude scaffolds largely consisting of unassembled repetitive regions or organelle sequence, we removed scaffolds < 100 kbp in length, scaffolds with fewer than one annotated gene per Mbp, and scaffolds with > 20% of the genes annotated as mitochondrial or chloroplast in origin. To assemble the mitochondrial and chloroplast genomes *de novo* from the reference genome’s PacBio circular consensus sequencing (CCS) reads we used PMAT v1.5.3^36^ to obtain a raw assembly graph in GFA format for each organelle. We then assessed coverage depth and manually resolved the graphs in Bandage v0.9.0^37^. The completed organelle genomes were added to the assembly, yielding a final reference of 18 chromosomes, two organelles, and 31 additional scaffolds.

### Generation of genomic sequencing data

We sequenced 90 novel specimens from Australian herbaria and generated additional sequencing data from previously constructed Illumina libraries for 127 North American and European herbarium samples described in Bieker *et al.*^11^ (Table S1). Leaf tissue samples were collected for DNA extraction. All pre-PCR steps were carried out in designated pre-PCR or clean room facilities. For herbarium specimens collected before 1960, DNA extraction was performed in a dedicated, positively pressurized, UV-irradiated ancient DNA laboratory at the NTNU University Museum, whereas samples collected after 1960 were extracted in the molecular genetics laboratories at the NTNU University Museum. DNA was extracted using the DNeasy Plant Mini kit (Qiagen) following the manufacturer’s instructions with an additional overnight incubation at 45 °C with 20 µL proteinase K after the lysis step as described in Martin *et al.*^38^. DNA concentration was quantified with a Qubit 2.0 fluorometer using the BR dsDNA kit and 2 µL extract. Extraction blanks were prepared alongside the samples to monitor for possible contamination. The distribution of fragment lengths of each DNA extract was assessed using a 2% agarose gel. If the mean fragment length was above ∼1,000 bp, the DNA was sheared before library preparation to a target mean fragment length of 500 bp using the Covaris 550-bp microTUBE-50 V2 program on a Covaris ME220 Focused-ultrasonicator (Table S1). For samples collected before 1960 that also required fragmentation, subsequent library preparation was carried out in the molecular genetics lab rather than the ancient DNA laboratory. Illumina libraries were prepared following the BEST library protocol^39^. Indexing PCR using dual indexing primers was performed as in Bieker *et al.*^11^, except that for each library two independent 50-µl PCR replicates were performed and then combined prior to purification. The optimal number of PCR cycles was assessed based on qPCR as described in Bieker *et al.*^11^. For the DNA purification after indexing PCR, Cytiva Sera-Meg beads were used with a 1:1 ratio for samples processed in the molecular lab and eluted in 33 µL EB buffer (Qiagen). The libraries were assessed for fragment size and molarity using High Sensitivity D1000 reagents and screentapes on the Agilent TapeStation 4200. Samples were pooled according to their estimated endogenous DNA proportion and sequenced on an Illumina NovaSeq X platform generating 150-bp paired-end reads.

### Alignment and SNP calling

We aligned resequencing reads spanning 790 samples^11,18^ (Table S2) to our reference using the RepAdapt pipeline^40^ (github.com/JimWhiting91/RepAdapt/tree/main/snp_calling_pipeline/mpileup_pipeline). Briefly, reads were trimmed and quality-filtered with fastp v0.20.0^41^ and aligned using bwa-mem v0.7.18 ^42^. Duplicates were removed using picard MarkDuplicates v2.21.4 (https://broadinstitute.github.io/picard/) and local realignment was performed around indels using GATK v3.8 IndelRealigner^43^.

To identify hybrids or misidentified species among the 90 new Australian herbarium samples, we first estimated genotype likelihoods from all herbarium-sample BAM files with ANGSD v.c877e7f^44^(`-SNP_pval 1e-6 -doMaf 2 -doGeno -1 -doPost 1 -minMapQ 25 - minQ 20 -trim 5 -minMaf 0.05 -geno_minDepth 2 -setMinDepthInd 2 - uniqueOnly 1`, with `-minInd` flag to 75% of the number of samples). We then pruned the genotype likelihoods with PLINK v.1.9 (`--indep-pairwise 50 5 0.5`), removed sites in genes and previously identified haploblock regions^18^, and generated a covariance matrix from 100,000 random SNPs drawn from the remaining sites in PCAngsd^45^. Eleven samples diverged strongly on PC1 and were excluded from further analysis.

To account for differences in sequencing quality between herbarium and non-herbarium (freshly collected tissue and silica-dried) samples, we called variants for each sample type separately using bcftools v.1.20^46^ `mpileup` and filtered with bcftools (`MQ < 30`) and vcftools v0.1.15 ^47^; `-- minQ 30 --minGQ 20 --minDP 3 --max-alleles 2 --max-missing 0.7`). We then removed ten samples with >60% missing data, and sites with mean depth <3X or more than 1 s.d. above the mean. Genotypes in the non-herbarium VCF were then phased and imputed with Beagle v5.1 ^48^, and the resulting imputed non-herbarium VCF was used as a reference panel to phase and impute variants in the herbarium VCF.

To assess residual differences between sample types, we performed discriminant analysis of principal components (DAPC) on the first 50 principal components to identify the axis separating herbarium and non-herbarium samples, and examined summary statistics for the SNPs loading most strongly on this axis. Because filtering more stringently on mapping quality most effectively shrank this axis, we repeated the filtering and imputation with a stricter mapping quality filter in bcftools, removing all sites with MQ < 59. This reduced the herbarium discriminant axis to a negligible fraction of the total variation in the PCA (0.08%) and indicated minimal systematic differentiation between herbarium and non-herbarium samples. The herbarium and non-herbarium VCFs were then joined. The final VCF contained 769 samples (405 non-herbarium; 364 herbarium) and 7,847,192 variant sites; after filtering for minor allele frequency (MAF) > 0.05, 1,589,190 sites remained.

### Climate data and derivation of bioclimatic variables

To exclude samples unsuitable for tracking allele frequencies across space and time, we removed samples missing collection location or year on the herbarium sheet. We then categorized samples as modern (collected in or after 1980) or historic (collected before 1980) and removed historic samples collected from beyond the geographic extent of the modern samples on each continent, leaving 735 samples (318 historic; 417 modern). We extracted climate data from the CRU TS v4.09 dataset (Climatic Research Unit, University of East Anglia), which provides globally gridded monthly climate variables at 0.5° resolution from 1901 to present. Monthly minimum temperature, maximum temperature, and precipitation were downloaded in decadal blocks spanning 1901–2020. For each ragweed sample, we used latitude, longitude, and sampling year to extract climate data over a nine-year window centred on the sampling year (year ± 4) using terra::extract in R. This window smooths interannual variability while capturing local climate conditions relevant to establishment and selection. Where sampling years fell near the temporal limits of CRU, the window was shifted to remain within the available range.

From the monthly data, we calculated 19 bioclimatic variables directly following standard WorldClim definitions, and removed variables that were highly correlated (|*r|* > 0.7) among sampling locations in any range, resulting in four minimally correlated bioclimatic variables for downstream analyses: BIO1 (mean annual temperature), BIO2 (mean diurnal temperature range), BIO12 (annual precipitation), and BIO15 (precipitation seasonality). To ensure comparability across ranges, environmental covariates were z-scaled using the mean and standard deviation of the native North American populations, and these scaling parameters were applied to all populations in all ranges.

### Genotype–environment association analyses

To identify climate-associated SNPs, we used the BayPass core model within each geographic range separately. Because BayPass requires population-level allele frequency estimates, analyses were restricted to modern samples, for which consistent population sampling was available. For each range we retained populations with >= 2 individuals per population (North America: 158 samples, 53 populations; Europe: 156 samples, 38 populations; Australia: 86 samples, 11 populations; Table S2) and filtered the resulting sites for MAF > 0.05. We included our four minimally correlated bioclimatic variables as environmental covariates and accounted for population structure via a range-specific Ω covariance matrix. To estimate Ω, we first generated a SNP set per range by LD-pruning in PLINK2 v.2.0-alpha (`--indep-pairwise 50 5 0.2`), excluding SNPs overlapping annotated genes (Battlay et al. 2023) and previously identified haploblock regions (Battlay et al. 2025), and randomly sampling 10,000 remaining SNPs. BayPass was then run under the core model to estimate Ω and allele-frequency prior parameters.

To estimate significance thresholds while accounting for demographic structure, we simulated a pseudo-observed dataset (POD) of 100,000 SNPs per range using the fitted Ω matrix and beta prior parameters, enforcing a minor allele frequency of 0.05. POD genotypes were reanalysed in BayPass under the same settings used for real data. For each environmental covariate, we extracted Bayes factor distributions from POD analyses and defined candidate SNPs as those exceeding the 99.9th percentile of the POD Bayes factor distribution for each covariate in each range.

To filter candidates to include only approximately independent loci, we estimated genome-wide linkage disequilibrium (LD) decay in each range separately using pairwise *r^2^* calculated from a random subset of 50,000 SNPs per range (MAF > 0.05). Pairwise *r^2^* values were computed in PLINK across distances up to 2 Mb (`--thin-count 50000 --r2 gz --ld-window 999999 --ld-window-kb 2000 --ld-window-r2 0`) and summarised as mean *r^2^* within overlapping 10-kb windows stepped every 1 kb in R. Across all ranges, mean *r^2^* declined rapidly within the first few tens of kilobases and approached background levels by ∼50 kb in each range, with consistently low LD (mean *r^2^*< 0.04) beyond this distance. Based on this decay profile, we clumped candidate SNPs using PLINK with a 50-kb window and an *r^2^* threshold of 0.1, retaining a single representative SNP per LD block for each range × environmental variable combination (`--clump-p1 1 --clump-p2 1 --clump-kb 50 --clump-r2 0.1`). These clumped candidate sets were used in all downstream analyses to reduce redundancy due to physical linkage while preserving signals of independent genotype–environment associations.

To generate null SNP sets matched to each clumped candidate set, we calculated modern allele frequencies per range and assigned each SNP to 5%-wide frequency tranches. For each tranche, we ensured null sets matched candidate sets in both SNP count and allele-frequency distribution by sampling the same number of SNPs from the same tranche without replacement after excluding haploblocks, genes, and the candidates themselves. These null sets were used as controls in all downstream analyses.

### Modelling spatial clines and temporal change

We modelled SNP-specific variation in population-level allele counts using generalized linear models implemented in *glmmTMB*^49^, fitted separately for each SNP, range, and environmental covariate. To control for population structure, we calculated principal components from PLINK2 LD-pruned (`--indep-pairwise 50 5 0.2`) genotype data within each range and included the first two axes (PC1 and PC2) in models as covariates. To evaluate alternative hypotheses about the tempo of adaptive change, time of sampling was modelled in three ways.

In all models, the response `cbind(a1_count, a2_count)` is the count of the two alleles per population, `env_c` is the mean-centred environmental variable, and `PC1_c` and `PC2_c` are the mean-centred population-structure axes. The models differ only in how sampling year enters the env × time interaction:

**Model 1 (continuous time):** cbind(a1_count, a2_count) ∼ env_c * year_c + PC1_c + PC2_c, where year_c is mean-centred collection year.

**Model 2 (two time categories):** cbind(a1_count, a2_count) ∼ env_c * histmod + PC1_c + PC2_c, where histmod classifies samples as historic or modern (pre- or post-1980).

**Model 3 (three time categories):** `cbind(a1_count, a2_count) ∼ env_c * histmod2 + PC1_c + PC2_c`, where `histmod2` classifies samples as early-historic (< 1900), mid- historic (1900–1980), or modern. Model 3 was fitted only in Europe, where temporal and spatial sampling were sufficient to split the historic samples of Model 2 into pre- and post-1900 groups of comparable sample size and temporal span.

All models were fitted using maximum-likelihood estimation with a betabinomial distribution (accounting for the overdispersion of allele counts) and logit link^50^. Model assumptions were checked using residual diagnostics in DHARMa^51^. Models failing to converge or violating assumptions were excluded from downstream analyses.

### Extraction of environmental cline slopes by time category

For categorical-time models (Models 2 and 3), we estimated environment-dependent cline slopes separately for each time category using *emmeans::emtrends()* with `specs = “histmod”` (or `histmod2`) and `var = “env_c”`, evaluated on the response scale. The resulting `env_c.trend` estimates correspond to allele-frequency cline slopes with respect to the environmental covariate in each time category. For each SNP, we recorded time-specific slope estimates together with standard errors and confidence intervals.

### Statistical analyses

All downstream analyses summarise the strength of spatial climate clines and their change through time using standardised effect sizes (estimate divided by standard error) derived from the genotype–environment–time models above. Temporal change was quantified in three complementary ways: *|Z_env×year_|*, the standardised environment × year interaction from the continuous-time model (Model 1); *|Z_env×histmod_|*, the standardised environment × time interaction from the two-category model (Model 2); and, in Europe where Model 3 was applied, *|ΔZ_env_|*, the absolute difference in standardised environmental effect size between consecutive time bins.

For analyses of asymmetry across environmental gradients, we evaluated the standardised temporal slope of allele frequency (the year slope divided by its standard error) conditional on environmental value, at the low (10th percentile) and high (90th percentile) ends of each gradient (*|Z_year|env_|*). For identifying climate-associated haploblocks in the native North American range (Fig. 6A) we used *|Z_env_|*, the environment effect from the continuous-time model (Model 1). Elsewhere (Figs 3, 4A), we used *|Z_env(mod)_|*, the environmental slope estimated at the modern time point in Model 2 (emtrends at `histmod = “modern”`).

To distinguish locus-specific from genome-wide change, climate-associated candidates were compared throughout against allele-frequency-matched null sets (above). Candidate and null distributions of a given statistic were compared with two-sided Wilcoxon rank-sum tests. *p*- values were reported per range × environmental variable without correction for multiple comparisons, as each addresses a distinct hypothesis. For analysis of parallelism in cline direction (Fig. 6B), the change in concordance between historic and modern samples was tested with McNemar’s tests.

### Locator source inference and climate mismatch

To infer the native-range source locations of historical samples from the invaded ranges, we used Locator, a supervised deep-learning method predicting geographic origin from genomic data^25^. Locator was trained using historical North American samples and their geographic coordinates, and predictions were generated for historical samples from the invaded ranges, Europe and Australia. To reduce the influence of any single genomic region on inferred source location, Locator was run separately on 10 Mbp non-overlapping windows spanning the genome. The final predicted source coordinates for each sample were calculated as the mean predicted longitude and latitude across all genomic windows.

We then compared the climate experienced by each historical sample at its observed collection location with the climate predicted for its inferred native-range source location. Inferred source locations did not show meaningful change over collection year in either invaded range (Fig. S5): Temporal trends explained <=2% of variance in source latitude and longitude (linear regression, all *r^2^* <= 0.021). Therefore all samples were assigned to a common historical time point corresponding to the median collection year of the historical dataset (1911) to make climate comparisons among historical samples comparable. Climate variables were then calculated from CRU data as above using a nine-year window centred on this year.

For each climate variable, we quantified climatic mismatch as the absolute difference between the standardized climate at a sample’s observed location and the standardized climate at its Locator-inferred native source location, with both values scaled to the mean and standard deviation of the native-range training samples (i.e. in native-range s.d. units). We tested whether mismatch was asymmetrically distributed across each environmental gradient by comparing mismatch at the lower and upper gradient extremes (25th vs 75th percentiles) using two-sided Wilcoxon rank-sum tests.

To assess whether inferred climate mismatch patterns were sensitive to uncertainty in Locator source assignment, we repeated the mismatch-gradient analysis using 1,000 bootstrap resamples of genomic windows. For each replicate, Locator windows were sampled with replacement, predicted source climate was recalculated as the mean across resampled windows, and mismatch-gradient models were refit.

### Flowering time genome-wide association and polygenic scores

We performed a genome-wide association study for onset of flowering using 212 modern samples from North America (*n* = 43), Europe (*n* = 85) and Australia (*n* = 84) for which phenotypic data had previously been measured by^19^. To account for relatedness and population structure, we LD-pruned SNPs in PLINK (`--indep-pairwise 50 5 0.2`), excluded SNPs overlapping annotated genes and haploblocks, and calculated an identity-by-state kinship matrix in PLINK (`--distance square ibs flat-missing`). The first two principal components calculated from the same LD-pruned SNP set were also extracted for use as covariates in the association model. GWAS was performed with EMMAX beta-07Mar2010 using the flowering onset phenotype and incorporating both the kinship matrix and PC covariates to control for relatedness and population structure among individuals. Genomic inflation was minimal (λ = 0.94; Fig. S6), indicating that relatedness and structure were adequately controlled. We used the flowering time GWAS results to calculate polygenic scores for all ragweed samples. To reduce redundancy among associated variants, GWAS SNPs were linkage disequilibrium clumped in PLINK using the 212 GWAS individuals, a 250 kbp window and an *r^2^* threshold of 0.2. We retained the lead SNP from each clump and generated four flowering time score sets using GWAS *p*-value thresholds of *p* < 1×10^-6^, *p* < 1×10^-5^, *p* < 1×10^-4^ and *p* < 1×10^-3^. For each threshold, individual polygenic scores were calculated in PLINK using the GWAS effect allele and estimated flowering time effect size for each retained lead SNP. Scores were calculated as the summed dosage of flowering time-increasing alleles weighted by their GWAS effect sizes, providing an individual-level estimate of genetic propensity for flowering time for all samples.

### Haploblock identification and genotyping

We identified haploblocks, extended genomic regions with distinct local population structure consistent with large structural variants, using local PCA^27^ across SNP data from all historical and modern samples. Haploblock identification was repeated using the combined historical and modern dataset to evaluate the persistence and distribution of previously identified structural variants and to identify any additional candidate haploblocks. Local PCA was performed in non-overlapping 10-kbp windows along each chromosome. For each chromosome, patterns of local population structure were summarised using multidimensional scaling, retaining 10 MDS axes. Candidate outlier regions were identified from the 5% corners of each pairwise combination of MDS axes and then manually inspected using chromosome-level MDS plots. Regions showing coherent clusters of outlier windows were retained as candidate haploblocks.

For each candidate region, we performed local PCA using SNPs within the candidate interval and calculated individual heterozygosity across the region. Haploblocks were genotyped from clustering patterns along PC1, PC2, or 45° rotations of these axes. We classified regions as haploblocks when samples formed three discrete clusters consistent with the expected genotypes of a large biallelic structural variant: two homozygous haplotype classes and a heterozygous class. Individual samples were assigned haploblock genotypes based on their cluster membership.

We validated haploblock genotype assignments using three complementary approaches. First, we tested whether samples assigned to the heterozygote cluster showed elevated heterozygosity across the candidate region relative to homozygote clusters using Wilcoxon tests, as expected for divergent haplotypes or inversion arrangements. Second, for chromosomes containing candidate haploblocks, we performed linkage disequilibrium scans comparing LD patterns in samples assigned to the common homozygote class with LD patterns in an *n*-matched random subset of samples. We considered loss or strong reduction of the LD block in the homozygote-only scan relative to the random subset as support for the haploblock genotype assignments. Finally, we compared candidate haploblocks to alignments between alternate haplotypes in the diploid reference genome^22^, providing independent support for candidate structural variants.

