## Supplementary figures S1-S6 for "Historical genomes reveal scale-dependent predictability of climate adaptation"

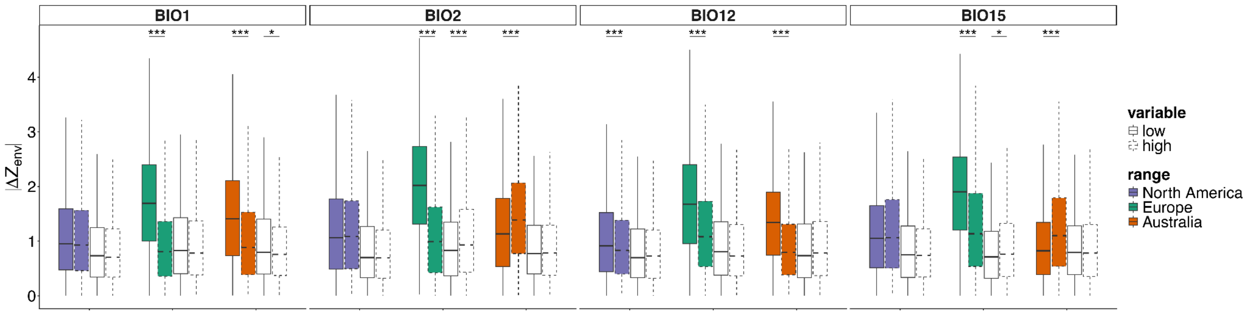


**Figure S1. Temporal cline shifts in range-specific candidate loci and frequency-matched nulls.** Distribution of temporal allele-frequency change at candidate loci (coloured boxes) and matched nulls (white boxes), quantified as the slope of allele frequency with respect to year, evaluated at the low (10th percentile) and high (90th percentile) values of each environmental gradient. Values are shown as standardized effect sizes (slope/SE; $|Z_{year|env}|$). Boxes show medians and interquartile ranges; whiskers indicate 1.5×IQR. Asterisks indicate significance of differences between groups in Wilcoxon tests (*$p < 0.05$; **$p < 0.01$; ***$p < 0.001$).

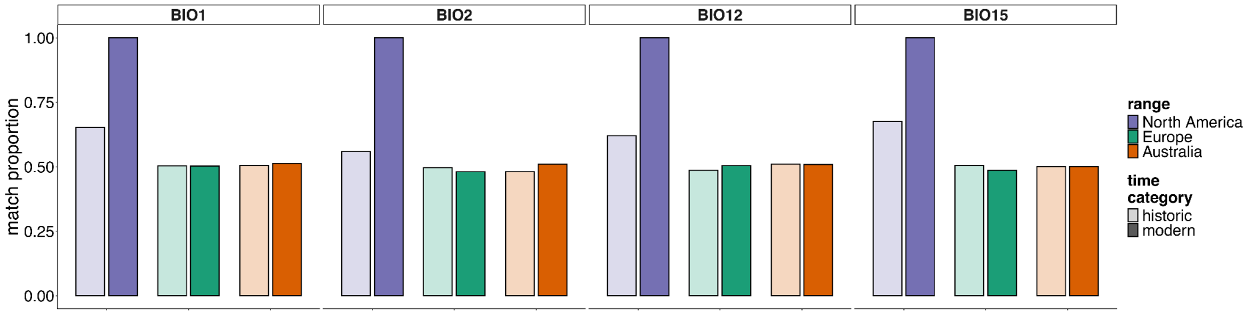


**Figure S2. Null-SNP directional alignment with native-range clines.** Bars indicate the proportion of null SNPs matched to North American candidate loci whose clines match the direction of the modern North American cline, shown separately for historic (pre-1980) and modern samples. Asterisks denote significant changes in matching frequencies between time categories based on McNemar's tests (*$p < 0.05$; **$p < 0.01$; ***$p < 0.001$). North American data are included for comparison, but McNemar's tests were not applicable because all loci matched the modern native cline by definition, resulting in degenerate contingency tables with no discordant pairs.

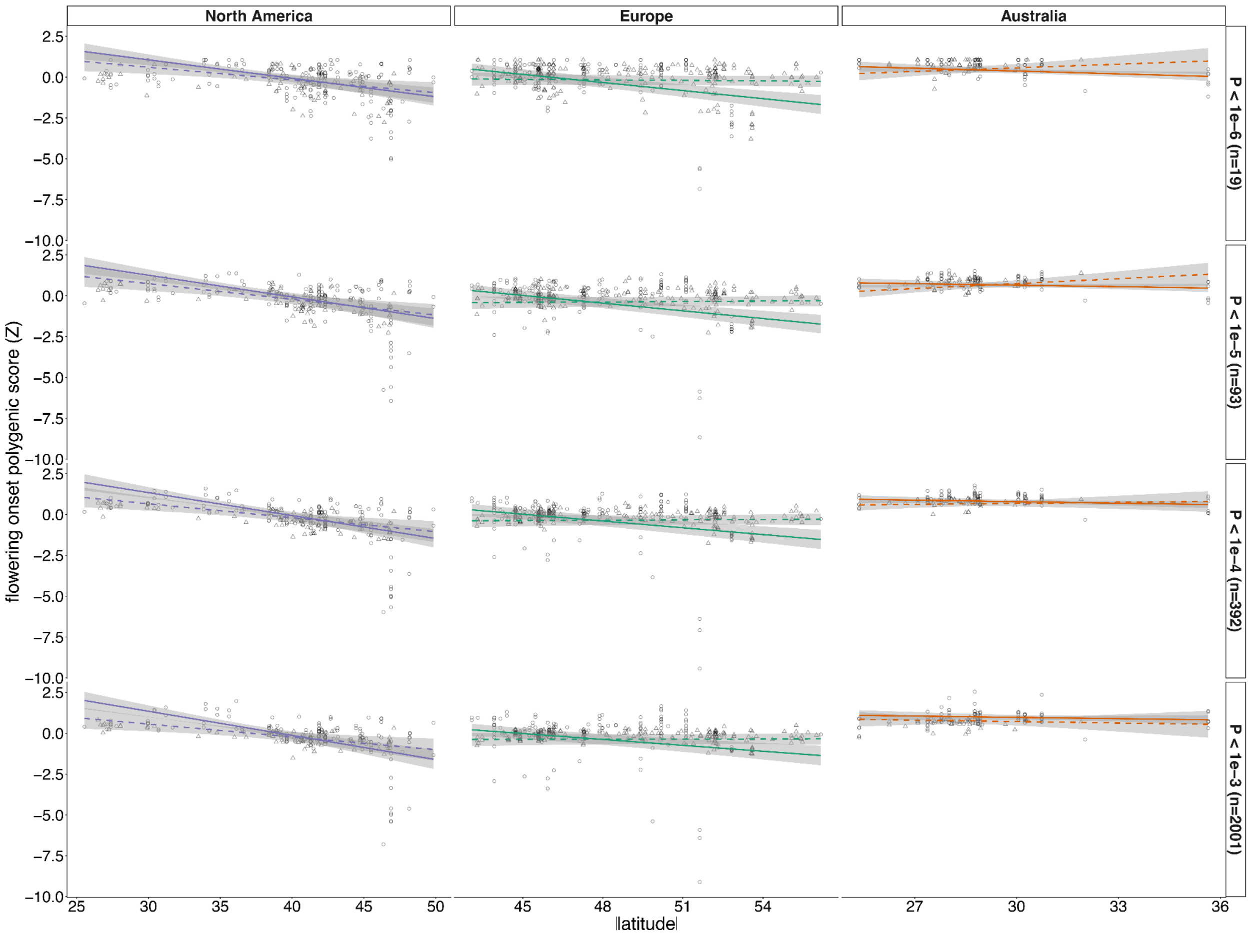


**Figure S3. Latitudinal clines in flowering onset polygenic scores across varying GWAS significance thresholds.** Polygenic scores (PGS; standardized) for flowering onset plotted against absolute latitude for each range (columns) for four GWAS significance thresholds (rows). Points represent individual samples (triangle = historic; circle = modern). Lines (dashed = historic; solid = modern) show fitted relationships from linear models including latitude, time (historic vs modern), and their interaction (with the first two principal components of genome-wide SNP variation as covariates); shaded regions indicate 95% confidence intervals.

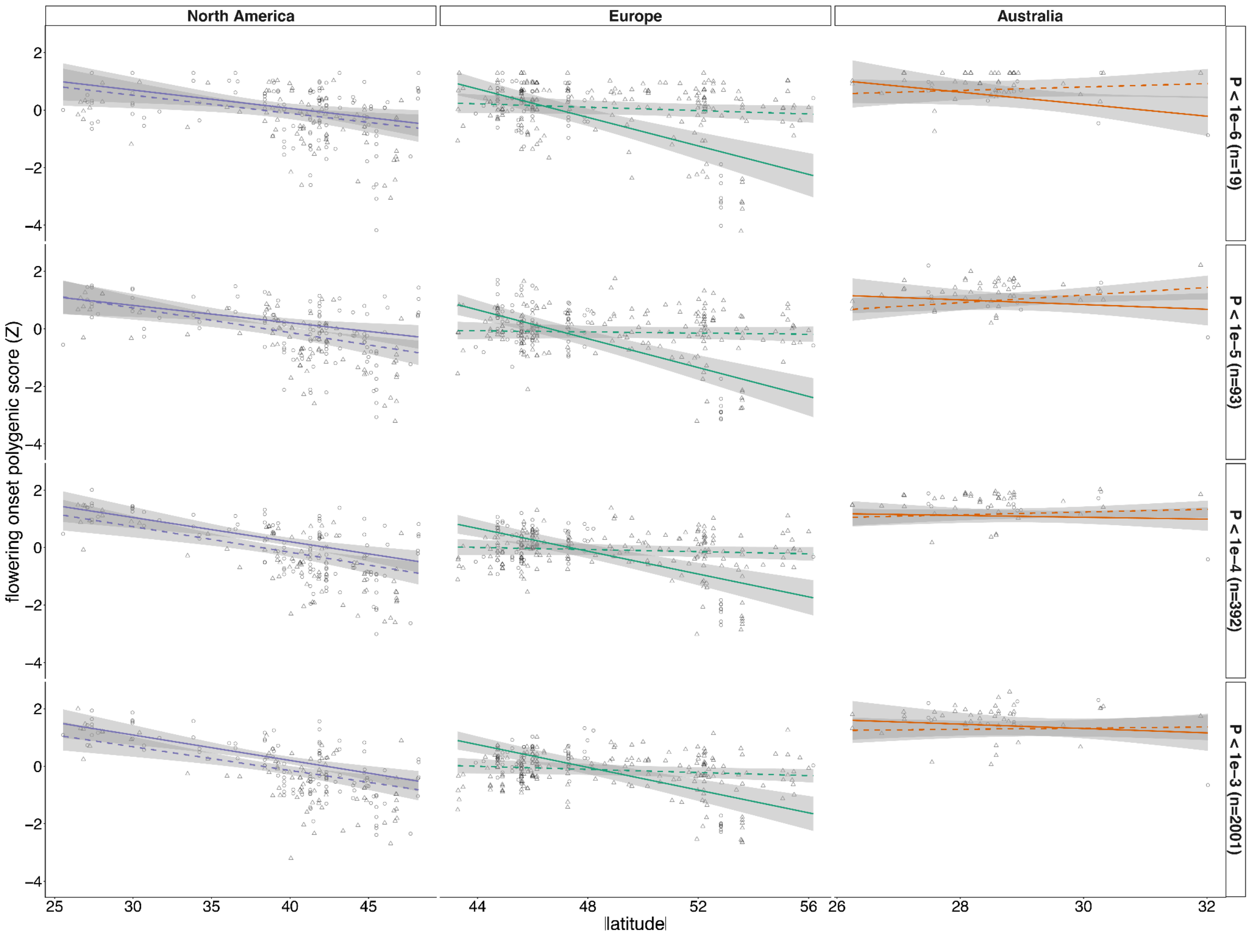


**Figure S4. Latitudinal clines in flowering onset polygenic scores across varying GWAS significance thresholds (GWAS samples excluded).** Polygenic scores (PGS; standardized) for flowering onset plotted against absolute latitude for each range (columns) for four GWAS significance thresholds (rows), with individuals used in the GWAS excluded from PGS prediction to guard against circularity. Points represent individual samples (triangle = historic; circle = modern). Lines (dashed = historic; solid = modern) show fitted relationships from linear models including latitude, time (historic vs modern), and their interaction (with the first two principal components of genome-wide SNP variation as covariates); shaded regions indicate 95% confidence intervals.

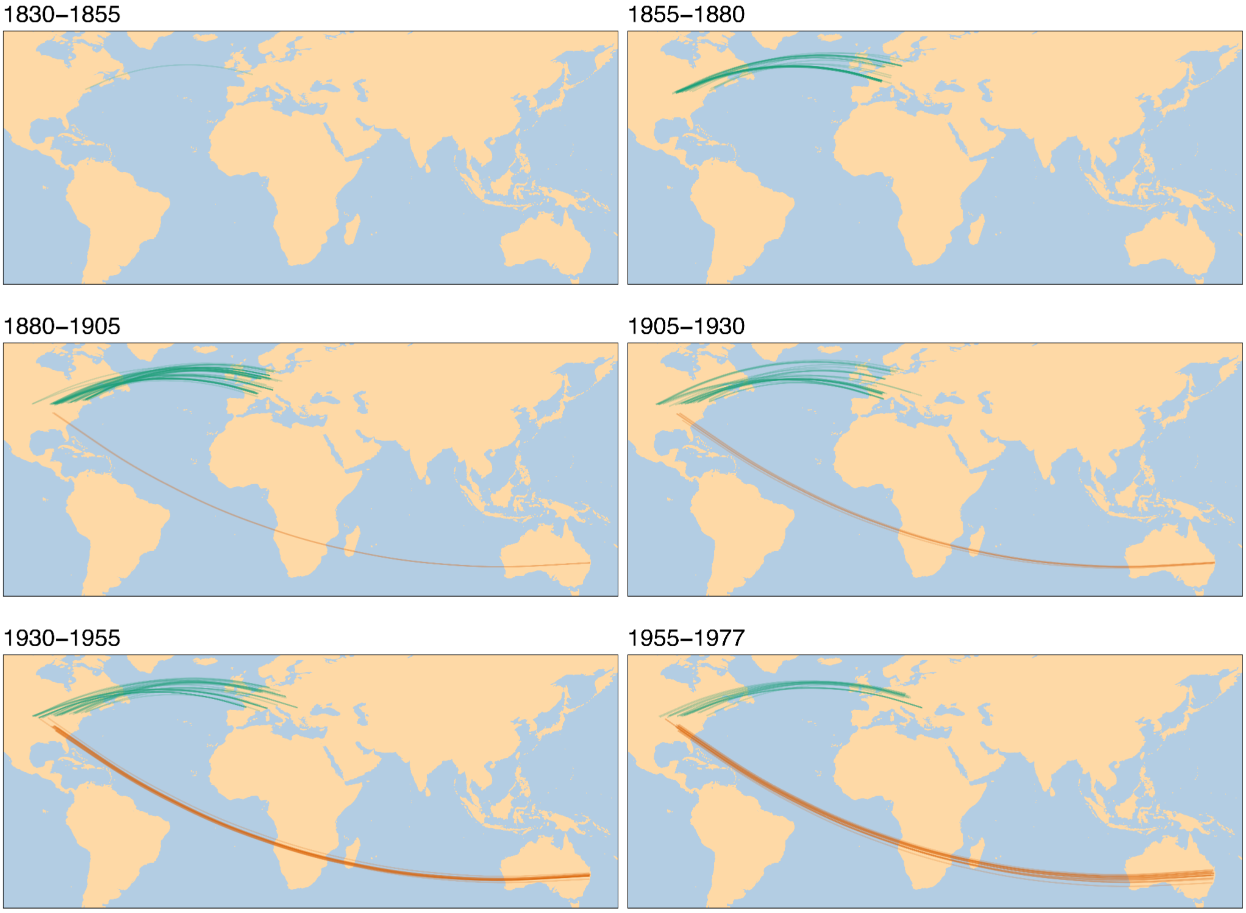


**Figure S5. Historic sample source locations over time.** Locator predictions for historic samples from Europe and Australia, based on models trained on historic North American samples. Lines connect sampling locations to their inferred source locations in the native range. Samples are divided into 25-year bins based on collection year.

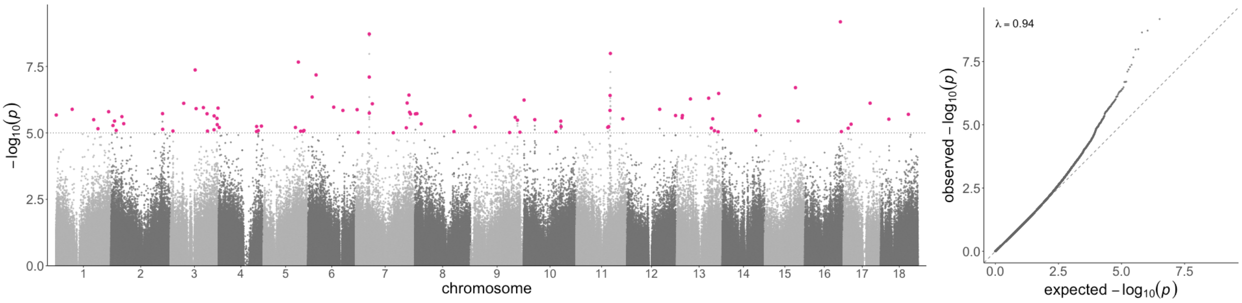


**Figure S6. Flowering-onset genome-wide association.** (left) Manhattan plot of $-\log_{10}(p)$ across the 18 chromosomes from an EMMAX mixed-model GWAS ($n$ = 212). The dotted line marks the $p$-value threshold used for polygenic-score construction ($p < 1\times10^{-5}$); pink points mark independent lead SNPs (LD-clumped, $p < 1\times10^{-5}$). (right) Quantile–quantile plot of observed versus expected $-\log_{10}(p)$; the genomic inflation factor (λ = 0.94) indicates well-controlled genomic inflation.
